# Determining insulin sensitivity from glucose tolerance tests in Iberian and Landrace pigs

**DOI:** 10.1101/2019.12.20.884056

**Authors:** JM Rodríguez-López, M Lachica, L González-Valero, I Fernández-Fígares

**Author notes:** This article has been peer-reviewed and recommended by *Peer Community in Animal Science* https://doi.org/10.24072/pci.animsci.100004.

## Abstract

As insulin sensitivity may help to explain divergences in growth and body composition between native and modern breeds, metabolic responses to glucose infusion were measured using an intra-arterial glucose tolerance test (IAGTT). Iberian (n = 4) and Landrace (n = 5) barrows (47.0 ± 1.2 kg BW), fitted with a permanent carotid artery catheter were injected with glucose (500 mg/kg BW) and blood samples collected at −10, 0, 5, 10, 15, 20, 25, 30, 45, 60, 90, 120 and 180 min following glucose infusion. Plasma samples were analysed for insulin, glucose, lactate, triglycerides, cholesterol, creatinine, albumin and urea. Insulin sensitivity indices were calculated and analyed. Mean plasma glucose, creatinine and cholesterol concentrations were lower (*P* < 0.01) in Iberian (14, 68 and 22%, respectively) compared with Landrace pigs during the IAGTT. However, mean plasma insulin, lactate, triglycerides and urea concentrations were greater (*P* < 0.001) in Iberian (50, 35, 18 and 23%, respectively) than in Landrace pigs. Iberian pigs had larger area under the curve (AUC) of insulin (*P* < 0.05) and lactate (*P* < 0.1), and smaller (*P* < 0.05) AUC for glucose 0-60 min compared with Landrace pigs. Indices for estimating insulin sensitivity in fasting conditions indicated improved β-cell function in Iberian compared with Landrace pigs, but no difference (*P* > 0.10) in calculated insulin sensitivity index was found after IAGTT between breeds. A time response (*P* < 0.05) was obtained for insulin, glucose and lactate so that maximum concentration was achieved 10 and 15 min post-infusion for insulin (Iberian and Landrace pigs, respectively), immediately post-infusion for glucose, and 20 min post-infusion for lactate, decreasing thereafter until basal levels. There was no time effect for the rest of metabolites evaluated. In conclusion, growing Iberian pigs challenged with an IAGTT showed changes in biochemical parameters and insulin response that may indicate an early stage of insulin resistance.

## Introduction

The Iberian pig is a slow growing native breed of the Mediterranean basin with much greater whole body fat content than lean-type pigs (Nieto *et al.*, 2002). Compared with conventional breeds, Iberian pigs show a lower efficiency of energy utilisation for protein deposition in the growing period (Barea *et al.*, 2007). The greater relative viscera weight (Rivera-Ferre *et al.*, 2005) and total heat production (González-Valero *et al.*, 2016) associated in part with the greater rate of muscle protein turnover (Rivera-Ferre *et al.*, 2005) in Iberian compared with lean-type pigs help to explain the low energy efficiency for growth. In fact, Rivera-Ferre *et al.* (2005) showed that muscle protein degradation was increased in Iberian pigs resulting in decreased muscle protein accretion compared with Landrace pigs. Interestingly, insulin resistance at the muscle level could explain an increased protein degradation (Wang *et al.*, 2006) affecting overall protein accretion. In a previous study using balanced or lysine deficient diets at two CP levels, Iberian had greater fasting serum insulin concentration than Landrace pigs (Fernández-Fígares *et al.*, 2007), suggesting the possibility of insulin resistance in Iberian pigs. We hypothesised that compared with modern lean breeds Iberian pigs have decreased insulin sensitivity, which could explain differences on growth, body composition and metabolic characteristics compared with modern breeds. The objective of the present study was to evaluate differences on insulin sensitivity between Iberian and Landrace pigs using an intra-arterial glucose tolerance test (IAGTT).

## Methods

### Animals

All procedures used in this study were approved by the Bioethical Committee of the Spanish National Research Council (CSIC, Spain), and the animals were cared for in accordance with the Royal Decree No. 1201/2005 (Spain). The experiment was performed with five Landrace and four Iberian (*Silvela* strain) barrows supplied by Granja El Arenal (Córdoba, Spain) and Sánchez Romero Carvajal (Jabugo S.A., Puerto de Santa María, Cádiz, Spain), respectively. The pigs were allowed *ad libitum* access to a standard diet (160 g CP/kg; 14 MJ metabolizable energy/kg DM) with free access to water in a controlled-environment room (21 ± 1.5°C). After acclimatization, each animal was surgically fitted with a chronic catheter in carotid artery following the procedure described previously (Rodríguez-López *et al.*, 2013). After surgery, the pigs were fed at 2.4 × metabolizable energy for maintenance (444 kJ/kg^0.75^ BW; National Research Council (NRC) 1998) with the same standard diet. On the day of the experiment, pigs (46.0 ± 3.0 and 47.8 ± 3.6 kg BW, Iberian and Landrace, respectively, that is about 18 weeks and 14 weeks, respectively) were given an intra-arterial bolus (500 mg/kg BW) of glucose (50% sterile dextrose; glucosado 50% Braun, B. Braun Medical S.A., Rubi, Barcelona, Spain) over one min period after an overnight fast. The catheter was immediately flushed with 5 mL of sterile saline solution. Blood samples (5 mL) were collected at −10, 0 (20-30 seconds after the bolus of glucose and the saline solution), 5, 10, 15, 20, 25, 30, 45, 60, 90, 120 and 180 min following glucose infusion.

### Biochemical analysis and calculations

Plasma was obtained by centrifugation (4°C, 1820 x g for 30 min; Eppendorf 5810 R, Hamburg, Germany) and stored in aliquots at −20°C until insulin and metabolites (glucose, lactate, triglycerides, cholesterol, creatinine, albumin and urea) were analysed. All samples were assayed in duplicate except for insulin which was assayed in triplicate.

Insulin was measured using commercially-available radioimmuno assay kit following the directions of the manufacturer (Millipore porcine insulin radioimmuno assay kit; Cat. PI-12K). Radioactivity in samples was measured using a gamma counter (Behring 1612; Nuclear Enterprises Ltd, Edinburgh, Scotland). Human insulin was used as standard, and the assay was validated for use in porcine plasma samples (Fernández-Fígares *et al.*, 2007). The intra- and inter-assay CVs for plasma insulin were 4.4 and 9.1%, respectively.

Plasma glucose, lactate, triglycerides, cholesterol, creatinine, albumin and urea were measured colorimetrically using an automated Cobas Integra 400^®^ analyser (Roche Diagnostics GmbH, Mannheim, Germany).

Responses of plasma insulin, glucose and lactate were evaluated separately by computing total area under the response curve (AUC) determined using trapezoidal geometry (GraphPad Prism, Version 5.02. San Diego, CA) for the time period indicated following intra-arterial glucose infusion (e.g. AUC0-5 stands for the integrated area between 0-5 min post-infusion, AUC0-10 between 0-10 min post-infusion, and so on, until AUC0-180). Basal levels per breed (at time −10 min) were used to calculate the corresponding AUC per metabolite. The rates of decline in plasma insulin and glucose concentrations for both breeds were calculated based on the slope in the linear portion of the response curve from 0 to 30 min after IAGTT challenge (Christoffersen *et al.*, 2009). Results were then expressed as a fractional rate constant determined from the slope of the natural logarithm of plasma concentrations vs. time (Shipley and Clark, 1972; cited by (Gopinath and Etherton, 1989). The fractional turnover rates (*k*), or disappearance rates, of plasma insulin and glucose (%/min) were calculated using the relationship (Kaneko *et al.*, 2008):

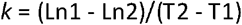

where Ln1 and Ln2 are the natural logarithms of plasma insulin concentration, (μU/mL, (or glucose concentration, mM) concentrations at times T1 (0 min) and T2 (30 min), respectively.

From the *k* value, the half-life, T_½_ (min), may be calculated as:

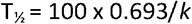

For insulin sensitivity, indices used in human medicine were used.

The so-called homeostasis model assessment (HOMA; Matthews *et al.*, 1985) was calculated for estimating insulin resistance (HOMA-IR) and β-cell function (HOMA-%B) at fasting conditions, as follows:

HOMA-IR: fasting plasma insulin (μU/mL) × fasting plasma glucose (mM)/22.5

HOMA-%B: (20 × fasting plasma insulin (μU/mL))/(fasting plasma glucose (mM) - 3.5)

It is assumed that non-insulin-resistant individuals have 100% β-cell function and an insulin resistance of 1.

The quantitative insulin sensitivity check index (QUICKI; Katz *et al.*, 2000) was computed as:

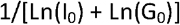

where I_0_ is the fasting insulin (μU/mL), and G_0_ is the fasting glucose (mg/dl).

Finally, the insulin sensitivity index (CSI; Tura *et al.*, 2010) was calculated as:

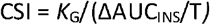

where *K*_G_ is the slope of Ln glucose in the linear portion of the response curve, ΔAUC_INS_ is the AUC of insulin above basal value, and T is the time interval (between 0 and 30 min) when *K*_G_ and ΔAUC_INS_ are calculated.

### Statistical analyses

The number of animals was calculated using the G*Power software (Heinrich-Heine-Universität Düsseldorf; (Faul *et al.*, 2007). Accepting an alpha risk of 0.05 and a beta risk of 0.2 in a two-sided test, **5** subjects are necessary in first group and **5** in the second to recognize as statistically significant a difference greater than or equal to 12 μU/ml on insulin concentration and a common standard deviation of 6.3 μU/ml based on previous studies (Fernández-Fígares *et al.*, 2007). A total of 5 pigs per treatment was also used by others (e.g. Stoll *et al.*, 1999). However, one Iberian pig lost the arterial catheter during the recovery period after surgery and only four Iberian pigs could be used.

Plasma metabolites were evaluated using a mixed ANOVA with repeated measures (Version 9.4; PROC MIXED, SAS Institute Inc., Cary, NC, USA) with the fixed effects of breed, time of sampling and their interaction in the model statement. The pig was considered a random effect. First-order ante dependence covariance ANTE(1) was used, which allows unequal variances over time and unequal correlations and covariance among different pairs of measurements. Plasma concentration differences between breeds at each sampling time were analysed by the pdiff (piecewise differentiable) option.

Assumptions that are required for an ANOVA were tested following the protocol from Zuur *et al.* (2010). Homogeneity of variance was assured by applying the Levene’s-Test. No transformation was required. Least square means and pooled SEM are presented. Differences were considered significant at *P* < 0.05 and trends approaching significance were considered for 0.05 < *P* < 0.10.

## Results

Fasting plasma insulin was greater in Iberian compared with Landrace pigs (15.6 *and* 8.10 μU/mL, respectively; sem=1.55, *P* < 0.05) whereas fasting plasma glucose was similar for both breeds (4.68 and 5.85 mmol/L for Iberian and Landrace pigs, respectively; sem=0.92, *P* > 0.10). No differences between breeds were found in fasting plasma albumin (Iberian 0.50 and Landrace 0.54 μM), urea (Iberian 3.3 and Landrace 3.0 mM), cholesterol (Iberian 1.46 and Landrace 1.79 mM) and triglycerides (Iberian 0.28 and Landrace 0.22 mM). On the other hand plasma fasting creatinine was lower in Iberian compared to Landrace (54 and 102 μM, respectively; *P* < 0.01). Average plasma metabolites and insulin concentrations after the IAGTT are shown in Table 1. Mean plasma glucose, cholesterol and creatinine concentrations were lower in Iberian (14, 22 and 68%, respectively; *P* < 0.05) compared with Landrace pigs. However, mean plasma insulin, lactate, triglycerides and urea concentrations were greater in Iberian (50, 35, 18 and 23%, respectively; 0.01 < *P* < 0.001) than in Landrace pigs. No differences (*P* > 0.1) were found between breeds for albumin levels.

**Table 1.** Average plasma metabolites and insulin concentrations in Iberian (n = 4) and Landrace (n = 5) pigs during an intra-arterial glucose challenge (IAGTT; 500 mg/kg BW, 0-180 min)^1^

|  | Breed |  | SEM | P-value<sup>2</sup> |  |  |
| --- | --- | --- | --- | --- | --- | --- |
|  | Iberian | Landrace |  | Breed | Time | Breed × Time |
| Insulin (μU/ml) | 41 | 27 | 1.9 | *** | *** | *** |
| Glucose (mmol/l) | 6.8 | 7.7 | 0.26 | ** | *** | ns |
| Lactate (mmol/l) | 1.3 | 1.0 | 0.039 | *** | *** | ns |
| Triglycerides (mmol/l) | 0.28 | 0.24 | 0.009 | ** | ns | ns |
| Cholesterol (mmol/l) | 1.5 | 1.8 | 0.033 | *** | ns | ns |
| Creatinine (μmol/l) | 54 | 90 | 1.2 | ** | ns | ns |
| Albumin (mmol/l) | 0.48 | 0.50 | 0.009 | ns | ns | ns |
| Urea (mmol/l) | 3.0 | 2.4 | 0.102 | *** | ns | ns |

<sup>1</sup>Average (n = 9) basal level of each metabolite: Glucose 5.33 mM, Insulin 11.43 μU/ml, Lactate 0.87 mM, Triglycerides 0.25 mM, Albumin 0.46 mM, Cholesterol 1.63 mM; Creatinine 76.2 μM, Urea 2.92 mM.
<sup>2</sup>ns = non-significant; \*\**P* < 0.01; \*\*\**P* < 0.001.

Only plasma insulin (Figure 1), glucose (Figure 2) and lactate (Figure 3) concentrations changed throughout time (*P* < 0.001; Table1) after the IAGTT.

**Figure 1.**
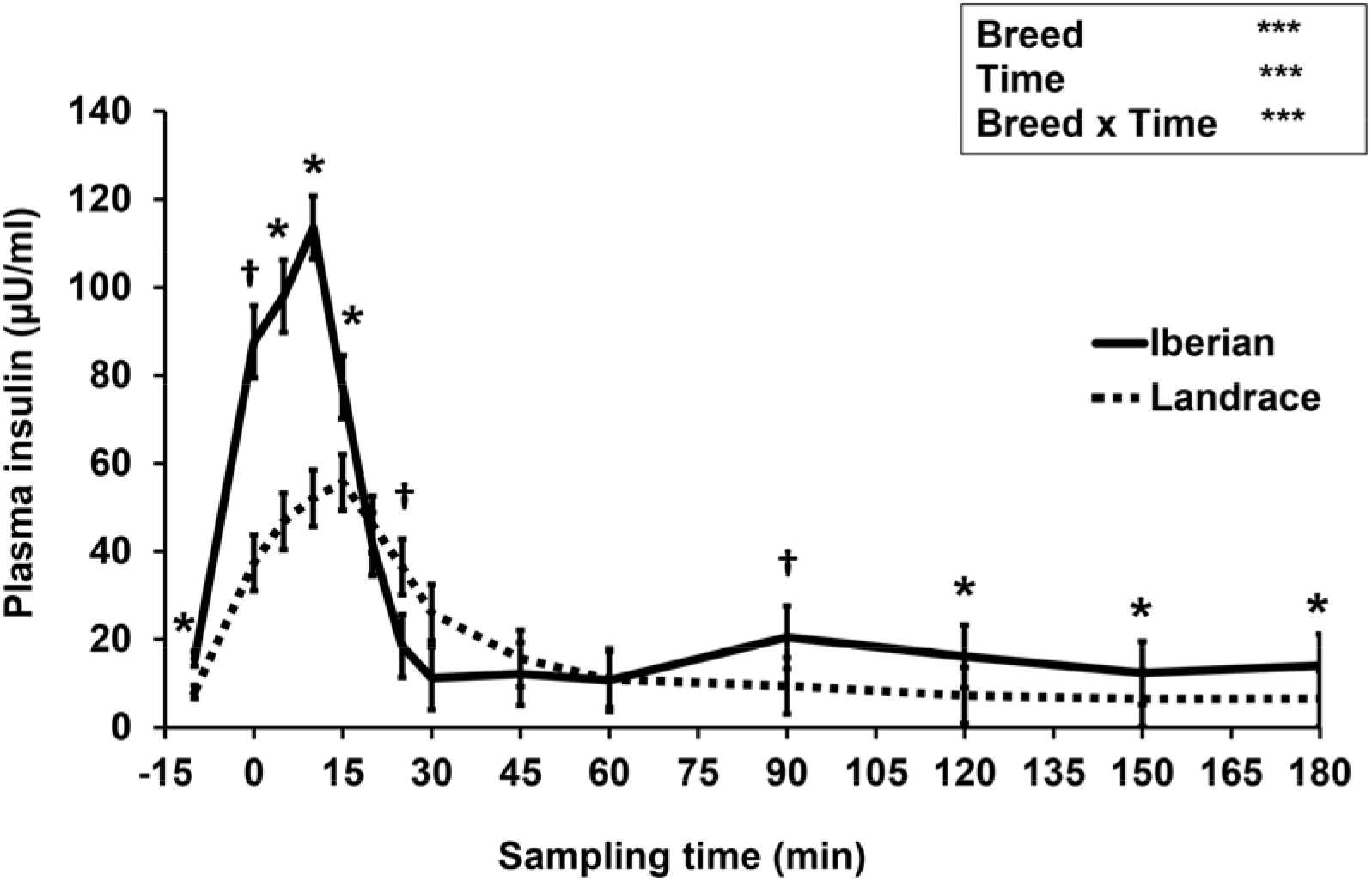
Plasma insulin concentrations during intra-arterial glucose challenge (500 mg/kg BW) in growing Iberian (n = 4) and Landrace (n = 5) pigs. Comparisons versus basal or control treatment: ^†^0.05 < *P* < 0.1, \**P* < 0.05, \*\*\**P* < 0.001. Average basal level of insulin 11.43 μU/ml.

**Figure 2.**
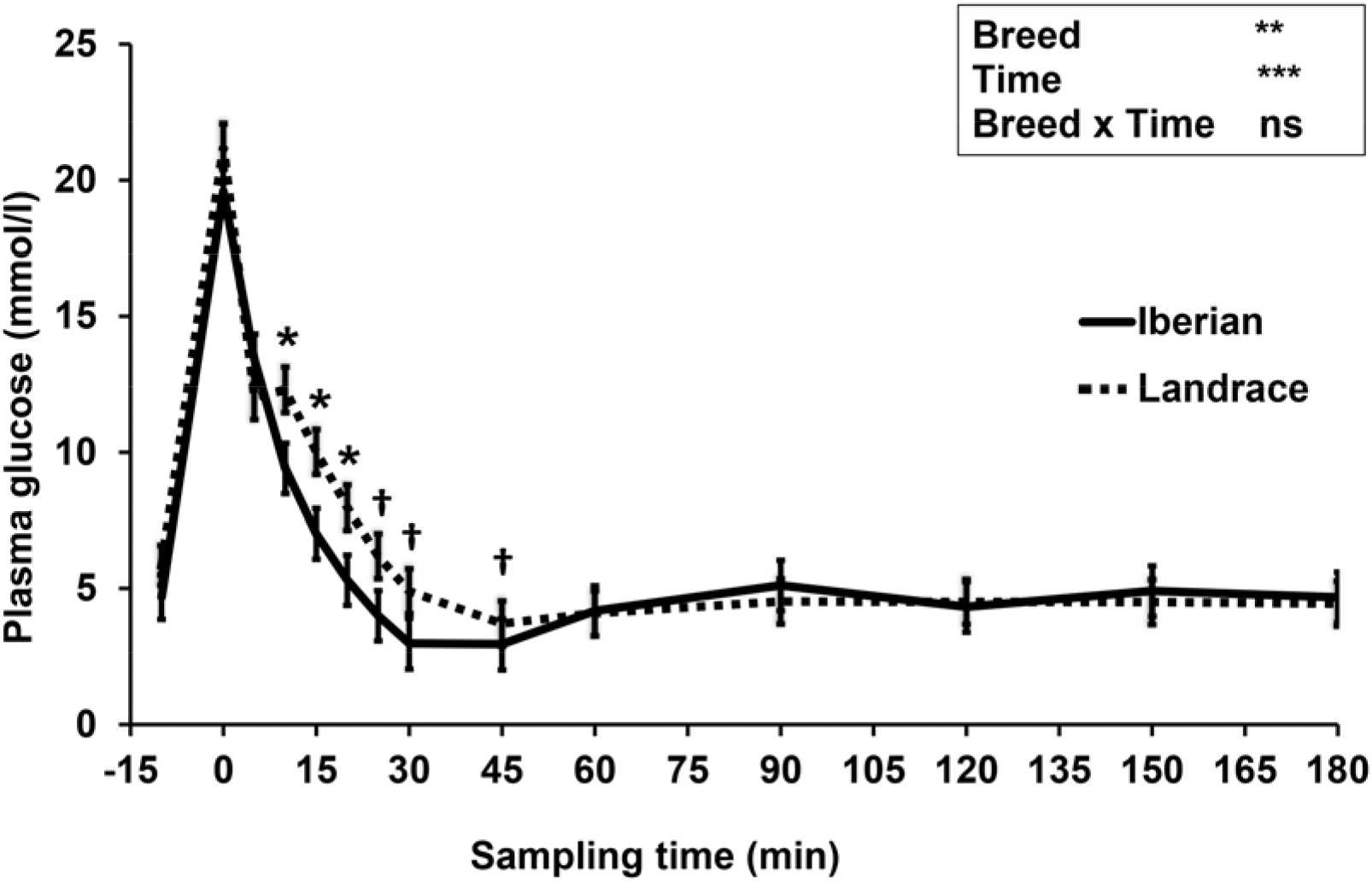
Plasma glucose concentration during intra-arterial glucose challenge (500 mg/kg BW) in growing Iberian (n = 4) and Landrace (n = 5) pigs. Comparisons versus basal or control treatment: ^†^0.05 < *P* < 0.1, \**P* < 0.05, \*\**P* < 0.01, *** *P* < 0.001; ns, not significant (*P* > 0.10). Average basal level of glucose 5.33 mM.

**Figure 3.**
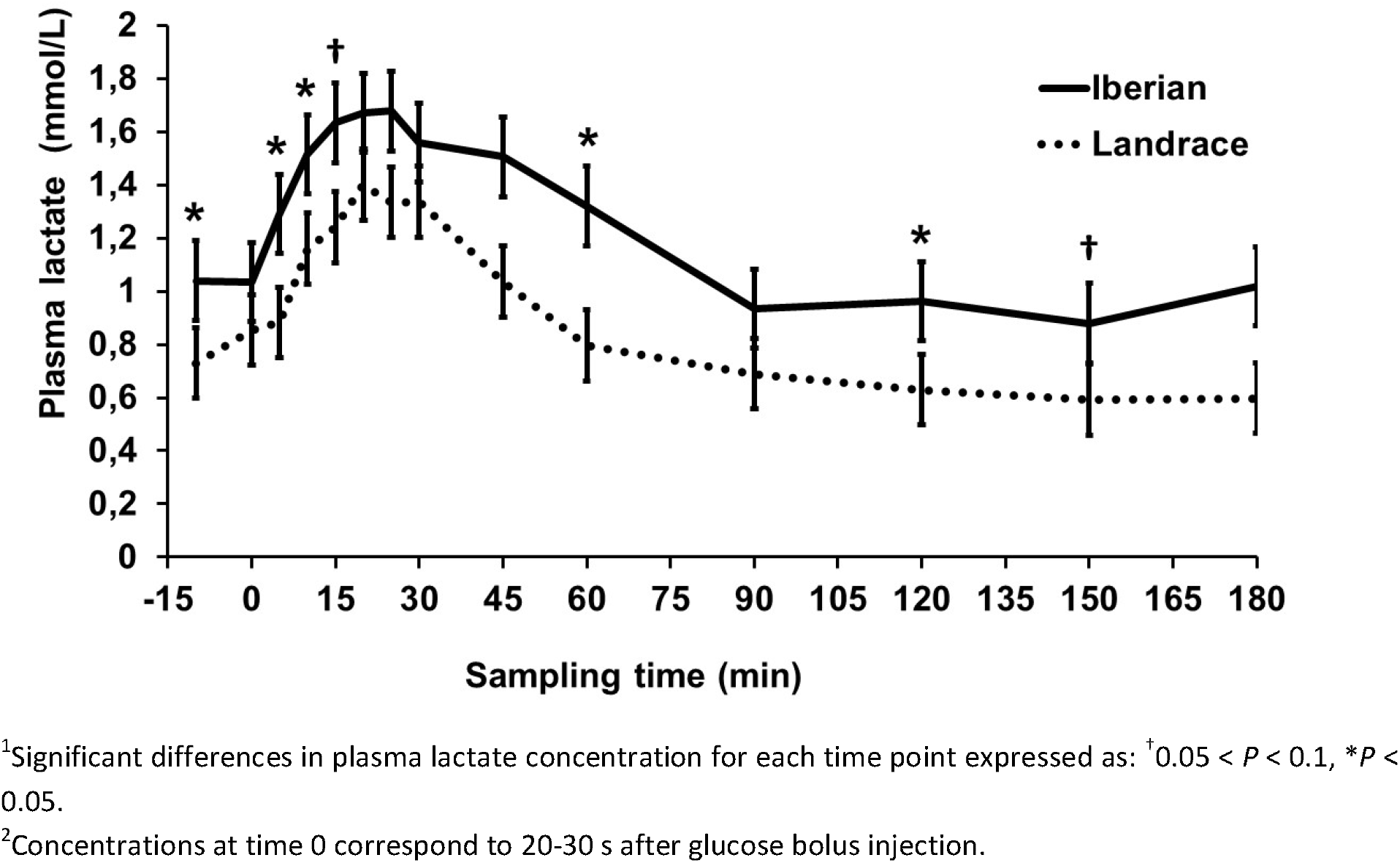
Plasma lactate concentration before and after intra-arterial glucose tolerance test (500 mg/kg BW) in growing Iberian (n = 4) and Landrace (n = 5) pigs^1, 2^.

An interaction between breed and time was found for plasma insulin, such that concentration of insulin was greater in Iberian pigs from −10 to 15 min and from 90-180 min (*P* < 0.05, with *P* < 0.1 at times 0 and 90 min) and lower at 25 min (*P* < 0.1; Figure 1). In both breeds, plasma insulin levels increased 7-fold, reaching a peak concentration 10 and 15 min after glucose infusion for Iberian (113.6 ± 7.1 μU/mL) and Landrace (55.7 ± 6.4 μU/mL) pigs, respectively. Insulin remained well above fasting levels until 20 and 45 min after glucose infusion for Iberian and Landrace pigs, respectively; thereafter insulin levels rapidly decreased until fasting levels were attained. Fractional turnover rate tended to increase in Iberian compared with Landrace pigs (6.2 and 4.7 ± 0.46%/min, respectively; 0.05 < *P* < 0.1) while half-life tended to decrease (11.3 and 15.2 ± 1.30 min for Iberian and Landrace pigs, respectively; 0.05 < *P* < 0.1).

Glucose peaked (Figure 2) immediately after glucose infusion reaching a value of 19.6 mmol/L and 21.2 mmol/L for Iberian and Landrace pigs, respectively. Subsequently, glucose concentration gradually decreased to values below fasting levels after 25 min and 30 min, respectively for Iberian and Landrace pigs. The lowest plasma glucose concentration (glucose nadir) was found at 45 min (2.95 mmol/L and 3.70 mmol/L for Iberian and Landrace pigs, respectively). After glucose nadir, glucose concentration gradually increased again to reach values comparable to fasting levels at 180 min. No differences were found between breeds for glucose fractional turnover rate (4.1 and 3.3 ± 0.37%/min for Iberian and Landrace pigs, respectively; *P* > 0.10) or glucose half-life (17.1 and 22.1 ± 2.67 min for Iberian and Landrace pigs, respectively; *P* > 0.10).

Lactate increased after the IAGTT, peaked at 20 min for both breeds and declined progressively until reaching basal concentrations at 180 min (Figure 3).

The AUC values for each sampling time of insulin, glucose and lactate are shown in Figure 4, 5 and 6, respectively. Insulin AUC was greater (*P* < 0.05) for Iberian compared with Landrace pigs at all times.

**Figure 4.**
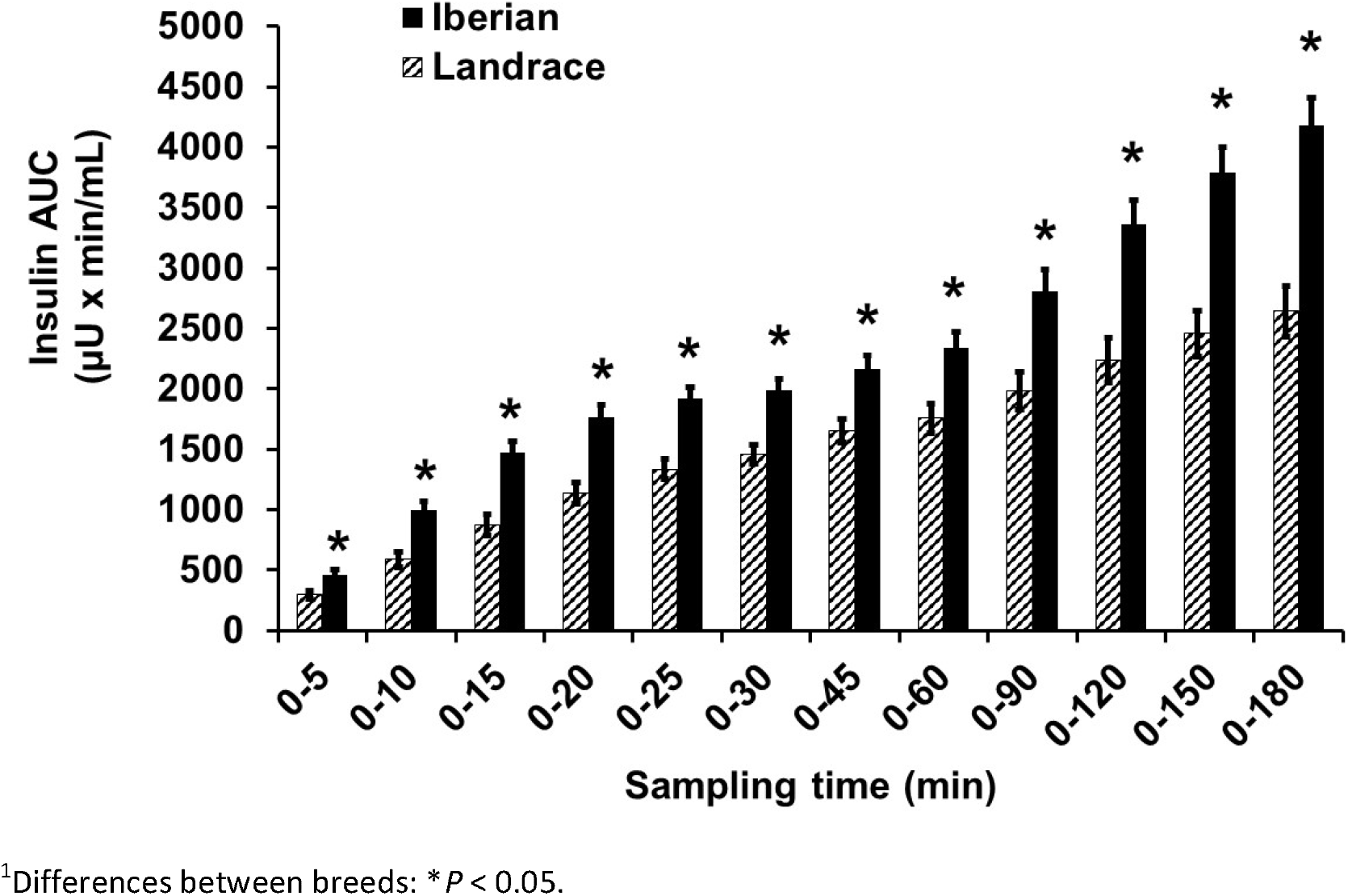
Area under the curve (AUC) of plasma insulin concentration during intra-arterial glucose tolerance test (500 mg/kg BW) between minute 0 and the time points indicated in growing Iberian (n = 4) and Landrace (n = 5) pigs^1^.

**Figure 5.**
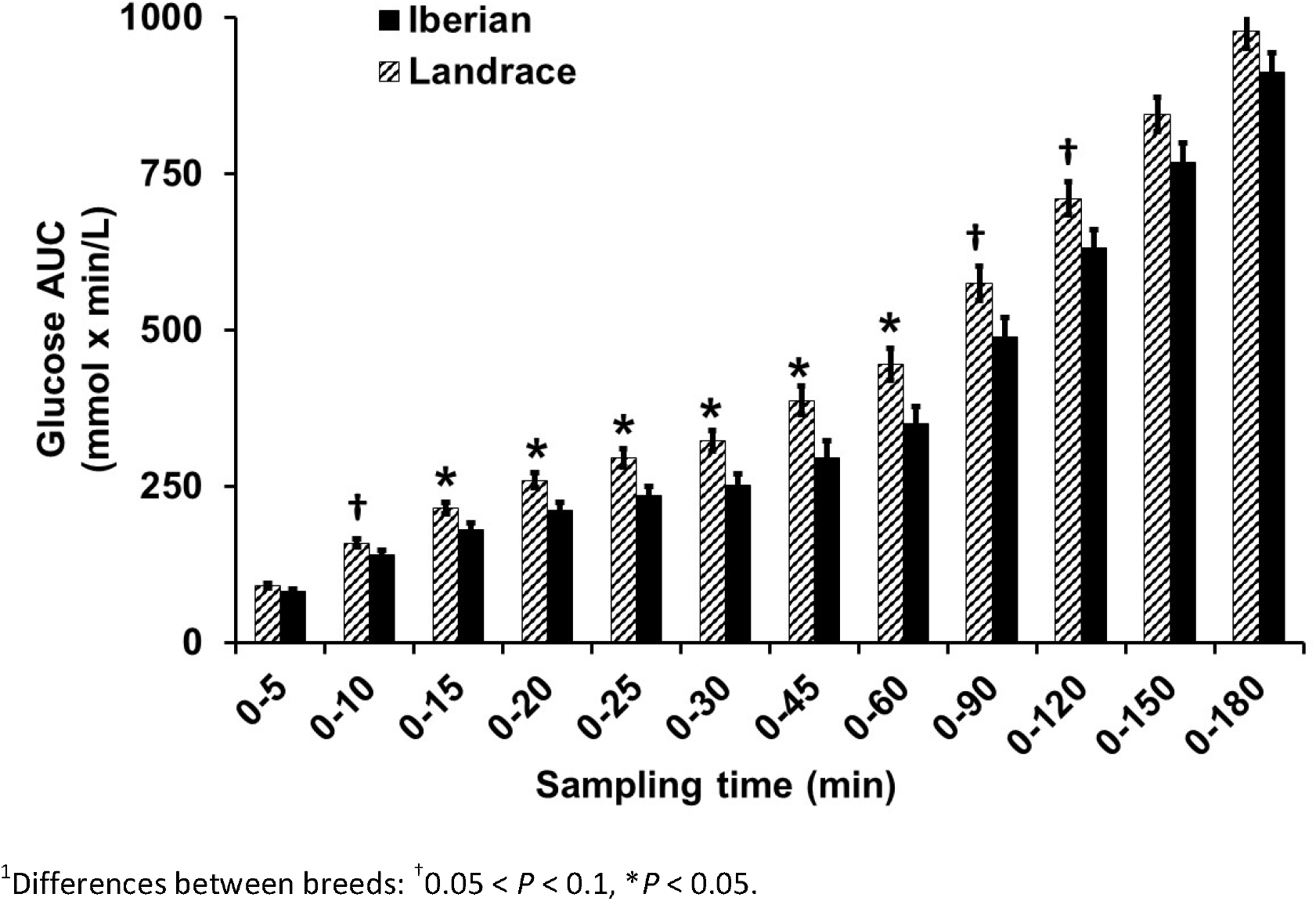
Areas under the curve (AUC) of plasma glucose concentration during intra-arterial glucose tolerance test (500 mg/kg BW) between minute 0 and the time points indicated in growing Iberian (n = 4) and Landrace (n = 5) pigs^1^.

**Figure 6.**
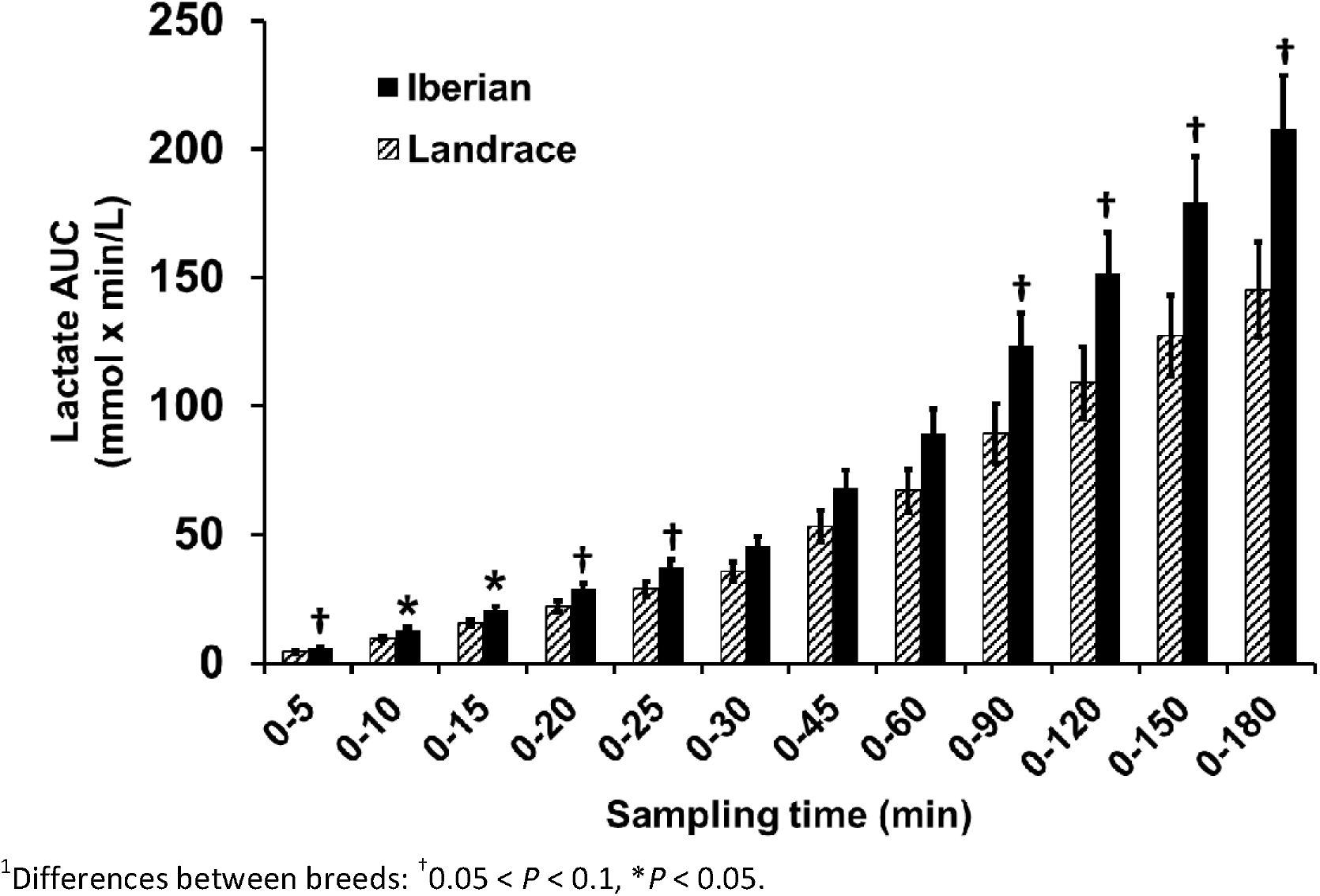
Area under the curve (AUC) of plasma lactate concentration during an intra-arterial glucose tolerance test (500 mg/kg BW) between minute 0 and the time points indicated in growing Iberian (n = 4) and Landrace (n = 5) pigs^1^.

Conversely, glucose AUC between 0-15, 0-20, 0-25, 0-30, 0-45 and 0-60 min were lower (*P* < 0.05) for Iberian than Landrace pigs. Glucose AUC tended to be lower (0.05 < *P* < 0.10) between 0-10, 0-90 and 0-120 min. No differences (*P* < 0.05) in glucose AUC was found between 0-5, 0-150 and 0-180 min. Plasma lactate AUC was greater (*P* < 0.05) for Iberian pigs for 0-10 and 0-15 min and tended to be greater (0.05 < *P* < 0.10) for 0-5, 0-20, 0-25, 0-90, 0-120, 0-150 and 0-180 min after glucose infusion.

Indices of insulin sensitivity are shown in Table 2. The QUICKI index decreased (*P* < 0.05) while HOMA-%B index increased (*P* < 0.01) in Iberian compared with Landrace pigs. No differences (*P* > 0.10) were found for HOMA-IR and CSI.

**Table 2.** Indices of glucose tolerance and insulin sensitivity in Iberian (n = 4) and Landrace (n = 5) pigs subjected to an intra-arterial glucose tolerance test^1^

|  | Iberian | Landrace | SEM | P-value<sup>2</sup> |
| --- | --- | --- | --- | --- |
| QUICKI | 0.31 | 0.33 | 0.007 | * |
| HOMA-IR | 3.3 | 2.3 | 0.58 | ns |
| HOMA-%B | 267 | 100 | 25.6 | ** |
| CSI (×10<sup>-4</sup>) | -12 | -13 | 1.8 | ns |

<sup>1</sup>QUICKI: quantitative insulin sensitivity check index; HOMA-IR: homeostasis model assessment for estimating insulin resistance; HOMA-%B: homeostasis model assessment for estimating β-cell function; CSI: calculated insulin sensitivity index.
<sup>2</sup>ns = non-significant; \*P < 0.05; \*\*P < 0.01.

## Discussion

The IAGTT method allowed the comparison of the insulin responsiveness of an obese (Iberian) and a lean (Landrace) pig breed. It has previously been shown that Iberian pigs are fatty pigs with slow growth rate compared to modern breeds (Nieto *et al.*, 2012). As arterial blood represents the metabolites concentration to which the tissues are exposed (Brouns *et al.*, 2005), chronic catheters were inserted in carotid artery for glucose infusion and blood sampling.

In the current study we have shown increased fasting plasma insulin and insulin response (higher plasma concentrations and AUC) after glucose infusion in Iberian compared with Landrace pigs (18 and 14 weeks of age, respectively). Greater postprandial serum levels of insulin have been described in 20 kg Iberian (11 weeks of age) compared to Landrace after glucose infusion (Fernández-Fígares *et al.*, 2007), and in 11kg Ossabaw (obese; 10 weeks of age) compared to 16.5 kg Yorkshire (10 weeks of age) pigs (Wangsness *et al.*, 1981). However, other comparative studies using a standard diet found increased insulin secretion in 75-120 kg Large White boars than in 40-75 kg Meishan boars (obese breed) at 20 and 52 weeks of age, respectively (Weiler *et al.*, 1998). The limited growth and development of slow growing pigs could result at least partly from disturbances in insulin secretion and/or in insulin binding, leading to insulin sensitivity, because most cells of the body require insulin for adequate uptake of glucose and amino acids (Claus and Weiler, 1994). If the concentration of insulin is compared among animals of different breeds, the sensitivity of each breed to insulin should be considered. In this study we were also interested in other key metabolites which could provide additional information concerning insulin sensitivity in Iberian pigs.

After glucose infusion, glucose plasma concentration rapidly returns to preprandial values as shown in the present experiment, which indicates that exogenous glucose was efficiently metabolized, stored as glycogen, or both. As expected, when glucose was infused, plasma glucose levels were rapidly increased and a subsequent insulin response was observed. The elevated insulin lowered plasma glucose below fasting values within 20 and 25 min for Iberian and Landrace pigs, respectively, and insulin levels returned to baseline as plasma glucose declined. In our study, glucose concentration and glucose AUC during the IAGTT were lower in Iberian compared with Landrace pigs, with no differences in fasting plasma glucose, maybe due to the limited number of pigs. When interpreting the individual glucose curves, a monophasic pattern was identified for both breeds. The lower glucose AUC of Iberian pigs (−19% on average) may be related to the greater insulin AUC (+33% on average), a common pattern in many models of obesity (Kay *et al.*, 2001). However, the reasons for the unequal physiological response between breeds are not well understood and must be discussed.

As it has been proved that the energy needs of portal-drained viscera are fulfilled by the oxidation of glucose, glutamate, and glutamine in pigs (Stoll *et al.*, 1999), a larger gastrointestinal tract of Iberian pigs compared to Landrace (Rivera-Ferre *et al.*, 2005) is in line with the decreased AUC of glucose reported in our experiment.

However, despite the larger size of the gastrointestinal tract and lower portal blood flow (González-Valero *et al.*, 2016) of Iberian compared with Landrace pigs, no differences on net portal flux of glucose after ingestion of the same diet were found (Rodríguez-López *et al.*, 2013). Differences on insulin stimulated glucose transport at portal-drained viscera level may help to explain these results. Iberian have lower glucose concentrations than Landrace pigs after an intravenous adrenaline challenge (Fernández-Fígares *et al.*, 2016), suggesting a decreased response of Iberian pigs to sympathetic nervous system stimuli which is in line with the lower glucose AUC reported here.

Lactate appearance after an intravenous glucose test is positively associated with insulin sensitivity in humans (Lovejoy *et al.*, 1992), as it is related to lactate production by insulin sensitive tissues (mainly muscle and fat). Because only limited amounts of lactate are produced by muscle after glucose loading (Ykijarvinen *et al.*, 1990), the source of lactate appearance should predominantly be adipose tissue (Lovejoy *et al.*, 1992), with a large capacity to convert glucose to lactate (Marin *et al.*, 1987). We report here a delay of 20 min in plasma lactate elevation relative to glucose peak following IAGTT, which may reflect the time lag in adipose tissue uptake of glucose and subsequent lactate production under the stimulation of insulin. Compared with Landrace, the increased lactate AUC in Iberian pigs after the IAGTT could therefore be a consequence of a larger adipose tissue (Nieto *et al.*, 2002) instead of greater insulin sensitivity. On the other hand, insulin resistance was associated with elevated basal lactate levels in obese humans (Lovejoy *et al.*, 1990), so increased basal lactate concentrations in Iberian pigs (1.040 vs. 0.730 mmol/L; sem=0.063) could also indicate insulin resistance or reduced insulin sensitivity. Although inhibition of insulin action on glycogenolysis in fasting conditions may lead to increased glucose release from glycogen and subsequent conversion of glucose to lactate, there is no direct evidence of this. There is indirect evidence, though, that elevated lactate levels reflect a glucose sparing effect (decreased glucose utilisation) in muscle (Pearce and Connett, 1980).

Obesity is frequently associated with different degrees of dyslipidemia manifested as increased triglyceridemia and low HDL-cholesterol. In our experiment, we found lower plasma cholesterol but greater plasma triglycerides concentration in Iberian compared with Landrace pigs, although we did not separate LDL and HDL fractions. Reduced total cholesterol concentration could be due to reduced hepatic insulin sensitivity as insulin stimulates cholesterol synthesis (Nelson and Cox, 2017). In any case the cholesterolemia for both breeds in the present experiment was in the lower range of published values (Fernández-Fígares *et al.*, 2007) and it cannot be considered that Landrace pigs were hypercholesterolemic.

Previous studies in our lab have shown the low genetic potential of growing Iberian pigs for muscle protein deposition in comparison to lean breeds (Nieto *et al.*, 2002), possibly due to the greater muscle protein degradation and turnover of the former (Rivera-Ferre *et al.*, 2005). In line with this, plasma urea level (an indirect protein degradation indicator) was in the present study 23% greater in Iberian compared with Landrace pigs. Differences on circulating insulin or the capacity of insulin release between breeds may explain differences in lean tissue deposition, as insulin has an important role in skeletal muscle metabolism (Wang *et al.*, 2006). In obese db/db mice (a model of insulin deficiency) higher muscle protein degradation in comparison with control mice (normal plasma insulin concentration) was reported; the authors concluded that insulin resistance was associated with accelerated muscle protein degradation (Wang *et al.*, 2006). The elevated protein degradation reported in Iberian compared with Landrace pigs (Rivera-Ferre *et al.*, 2005) suggests the possibility of insulin resistance at this level. The lower plasma creatinine level (indicator of muscle mass) found in this study for Iberian pigs, is in accordance with previous studies (Fernández-Fígares *et al.*, 2007) and also with the low muscle protein deposition and muscle size described previously (Nieto *et al.*, 2002; Rivera-Ferre *et al.*, 2005). As insulin resistance is associated with decreased muscle mass, plasma creatinine levels can also be used as an indicator of insulin signalling disorders as reported by Kashima *et al.* (2017) in humans. Further research regarding amino acid concentration after an IAGTT may help to explain differences in the effect of insulin on muscle protein metabolism between breeds.

When insulin sensitivity indices used in human medicine were applied to the conditions of the present experiment, QUICKI and HOMA-%B were more sensitive detecting differences between breeds. Indeed, QUICKI index decreased in Iberian compared with Landrace pigs, pointing out an incipient insulin sensitivity impairment in fasting Iberian pigs. Similarly, reduced QUICKI index (0.5 vs. 0.6) was found in Bama miniature pigs fed a high sucrose high fat diet compared with a control diet, respectively (Liu *et al.*, 2017). The QUICKI index has been shown to provide reasonable approximations of insulin efficiency in minipigs (Christoffersen *et al.*, 2009).

When we used the homeostasis model assessment (HOMA), differences on hepatic insulin resistance (HOMA-IR index) were negligible between breeds (3.3 and 2.3 for Iberian and Landrace, respectively; P>0.10). However, Iberian had improved β-cell function compared with Landrace pigs according to HOMA-%B index (267 and 100 for Iberian and Landrace, respectively; P<0.01), which may be due to enhanced sensitivity of the β-cells to glucose during the fasting period. As a consequence, β-cell insulin synthesis in Iberian pigs increased in accordance with the increased insulin release after the glucose tolerance test and the elevated basal insulin concentrations reported for Iberian pigs. This is consistent with decreased quantitative insulin sensitivity check index in Iberian pigs compared to Landrace (0.31 and 0.33 for Iberian and Landrace, respectively; P<0.05).

Previous studies from our lab indicate that growing Iberian pigs are prone to insulin resistance compared with modern breeds as denoted by increased hepatic gluconeogenesis (González-Valero *et al.*, 2014), greater plasma free fatty acid concentration (Fernández-Fígares *et al.*, 2016) and lower plasma creatinine and QUICKI index (Fernández-Fígares *et al.*, 2007). Additionally, in this experiment we show greater HOMA-%B and increased plasma insulin and lactate concentrations after an IAGTT. The increased plasma insulin AUC after an IAGTT suggests insulin resistance in comparison to values obtained for lean pigs, although the concentration of glucose remained low which could indicate the absence of a peripheral insulin resistance. Although Iberian pigs may be considered an obese breed in terms of body composition (Nieto *et al.*, 2002; Barea *et al.*, 2007), insulin resistance mechanisms have not yet been fully established at the development stage of the pigs in this experiment. Insulin resistance and impaired glucose tolerance has been shown in Iberian sows (2.5 years old) *ad libitum* fed a saturated fat enriched diet for three months (Torres-Rovira *et al.*, 2012). Although our results support the existence of an insulin resistance or a decreased insulin sensitivity in growing Iberian pigs, caution should be taken because of the reduced number of pigs used. The utilization of the hyperinsulinemic euglycemic clamp, the most definitive approach to determine whole-body insulin action should provide conclusive evidence regarding the establishment of insulin resistance in growing Iberian pigs.

## Data accessibility

Data are available online: https://doi.org/10.5281/zenodo.3609520

## Supplementary material

There is no supplementary material.

## Acknowledgements

The authors thank the company Sánchez Romero Carvajal (Jabugo S.A., Puerto de Santa María, Spain) for their helpful collaboration, Dr Luis Lara for statistical advice and Dr Thomas J. Caperna for critically reading the manuscript. This research was supported by grant AGL2006-5937 from the Ministry of Science and Education of Spain. Version 3 of this preprint has been peer-reviewed and recommended by Peer Community In Animal Science (https://doi.org/10.24072/pci.animsci.100004).

## Appendix

No appendix is provided.

